# Recombinant SARS-CoV-2 genomes are currently circulating at low levels

**DOI:** 10.1101/2020.08.05.238386

**Authors:** David VanInsberghe, Andrew S. Neish, Anice C. Lowen, Katia Koelle

## Abstract

Viral recombination can generate novel genotypes with unique phenotypic characteristics, including transmissibility and virulence. Although the capacity for recombination among betacoronaviruses is well documented, there is limited evidence of recombination between SARS-CoV-2 strains. By identifying the mutations that primarily determine SARS-CoV-2 clade structure, we developed a lightweight approach for detecting recombinant genomes. Among the over 537,000 genomes queried, we detect 1175 putative recombinants that contain multiple mutational markers from distinct clades. Additional phylogenetic analysis and the observed co-circulation of predicted parent clades in the geographic regions of exposure further support the feasibility of recombination in these detected cases. An analysis of these detected cases did not reveal any evidence for recombination hotspots in the SARS-CoV-2 genome. Although most recombinant genotypes were detected a limited number of times, at least two recombinants are now widely transmitted. Recombinant genomes were also found to contain substitutions of concern for elevated transmissibility and lower vaccine efficacy, including D614G, N501Y, E484K, and L452R. Adjusting for an unequal probability of detecting recombinants derived from different parent clades, and for geographic variation in clade abundance, we estimate that at most 5% of circulating viruses in the USA and UK are recombinant. While the phenotypic characterization of detected recombinants was beyond the scope of our analysis, the identification of transmitted recombinants involving substitutions of concern underscores the need to sustain efforts to monitor the emergence of new genotypes generated through recombination.

## Introduction

Severe Acute Respiratory Syndrome Coronavirus 2 (SARS-CoV-2) emerged in December of 2019 in China but has since spread worldwide. Laboratories around the world have been sequencing and rapidly sharing SARS-CoV-2 genomes throughout the pandemic, providing researchers the rare opportunity to study the evolution of SARS-CoV-2 in real-time. As of February 16, 2021, 537360 complete viral genomes and 300592 unique genotypes were available on the online repository GISAID^1^.

In addition to point mutations and insertions/deletions, coronavirus evolution is heavily driven by recombination^2^. Recombination events create chimeric genotypes between two viral strains that infect the same cell. This process occurs when RNA polymerase prematurely stops replicating the first genotype before reassembling and resuming replication with the second genotype as template. The end result is the unlinking of mutations across the genome, creating novel combinations of existing mutations. The clinical and epidemiological relevance of these new combinations is substantial as they have the potential to create genotypes with unique virulence and transmissibility characteristics.

Measurements of the frequency of recombination among coronaviruses in cell culture suggest it is very common^3–5^. There have further been attempts to detect and measure the magnitude of recombination among naturally circulating SARS-CoV-2 genomes. Based on four single nucleotide polymorphisms (SNPs), an early analysis reported recombinants among the first 85 sequenced SARS-CoV-2 genomes^6^. Two more recent pre-prints have also identified recombinant SARS-CoV-2 genomes, but used substantially different methods^7,8^. Although these analyses identified recombinant SARS-CoV-2 genomes, four studies have reported evidence of strong linkage disequilibrium among polymorphic sites and no disruption of the clonal pattern of inheritance, suggesting recombinant SARS-CoV-2 strains are not widespread^9–12^.

Here we add to these existing studies by adopting a lightweight approach to rapidly screen for recombinant SARS-CoV-2 genomes. Using this approach, we identify over 300 unique recombinant genomes and use phylogenetic placement to demonstrate statistical support for recombination. Accounting for the uneven phylogenetic distances between circulating viral genotypes and changes in their prevalence over time, we estimate that at most 5% of circulating strains in the United Kingdom and USA are recombinants. Finally, to facilitate future screening efforts, we implemented a local alignment-based version of our recombination detection approach that can rapidly screen large genome databases with limited resources.

## Results

### The limited genome-wide diversity among SARS-CoV-2 strains restricts the ability to detect recombinants

Compared to RNA viruses that have been endemically circulating in humans, SARS-CoV-2 harbors only a small amount of genetic variation. Of the approximately 30 thousand sites in the SARS-CoV-2 genome, only 121 positions have nucleotide variants that are present in at least 1% of genomes sequenced to date, and only 15 positions have variants present in at least 10% of genomes. Consequently, the clade structure of SARS-CoV-2 is overwhelmingly determined by a small number of variant sites. An efficient approach for identifying recombinant SARS-CoV-2 genomes could therefore rely on detecting unusual combinations of single nucleotide polymorphisms (SNPs) which are phylogenetically informative. This approach has the added benefit of not being computationally demanding, which facilitates the process of screening large sequence databases.

To define phylogenetically informative variant sites, we identified SNPs that are strongly associated with major phylogenetic clades within SARS-CoV-2. Multiple nomenclature schemes have been introduced to describe the major lineages that are currently in circulation, the most widely used of which are the Nextstrain^13^, GISAID^14^, and Pangolin^15^systems. While Pangolin provides a fine-scaled classification of viral diversity, Nextstrain and GISAID provide a broad-scale classification scheme, clustering genomes into five or seven clades, respectively. To increase the number of variant sites that contribute to our detection of recombinant genomes, we increased the resolution of the Nextstrain scheme by further dividing their clades into 14 monophyletic clades with at least 50% bootstrap support (Fig. 1A). We then screened all polymorphic sites in a genome alignment of over 6000 high quality genome sequences to identify SNPs that are strongly associated with the 14 clades. These clade-defining SNPs (cdSNPs) are defined as sites where >95% of the members in at least one clade have an alternative nucleotide to the dominant allele at that site, and that same alternative nucleotide is present in <5% of the strains in all remaining clades. For example, 99.5% of the members of clades 20B-1 have a T at site 23731 while the remaining 13 clades have a C at that same site, such that site 23731 is included as a cdSNP (Fig. 1C). In total, we identified 37 cdSNPs that reliably distinguish all 14 clades (Fig. 1 ABC). Among these positions is the nucleotide substitution of A to C at site 23403, which is responsible for the D614G mutation in the spike protein that has been associated with increased transmissibility^16–18^. Of the over half a million complete, human-derived SARS-CoV-2 strains available on GISAID as of February 16, 2021, fewer than 0.5% of genomes differed from the 14 clades’ cdSNP profiles by more than one nucleotide.

**Figure 1.**
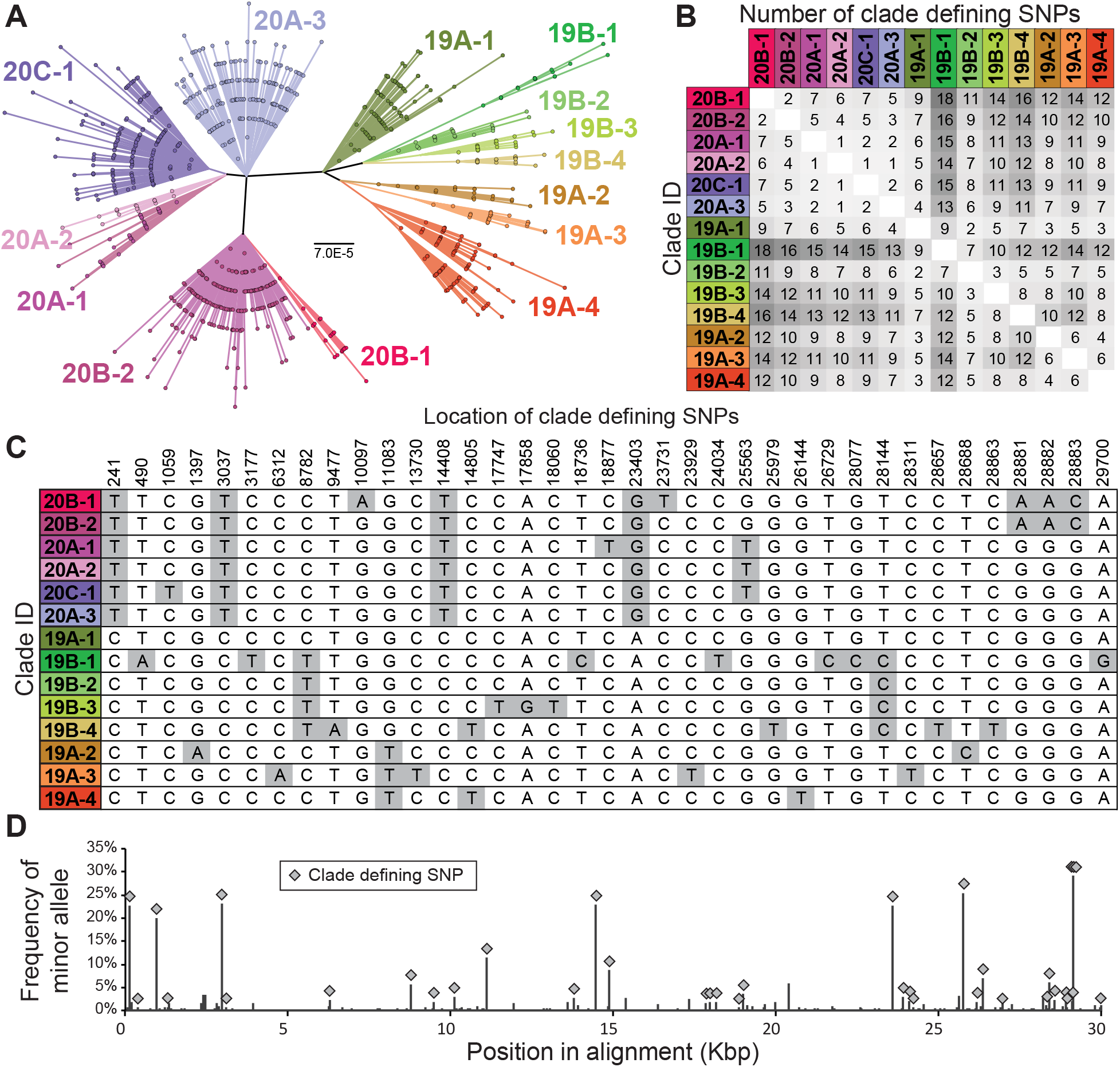
The clade structure of SARS-CoV-2 is shaped predominantly by 37 clade-defining SNPs. **(A)** An unrooted, maximum likelihood phylogeny based on the General Time Reversible model with invariant sites of 6536 high quality unique genome sequences with <1% Ns. The high-quality sequences used spanned collection dates between January 22 and May 17, 2020. 14 monophyletic clades with ≥50% bootstrap support were identified and named based on Nextstrain nomenclature19. Clade boundaries are in full agreement with Nextstrain clades (e.g., 20B), but some are more finely differentiated for higher resolution in our analyses (e.g., 20B-1). Scale bar is in substitutions per site. **(B)** Pairwise differences between the clade-defining SNP profiles of all 14 clades. **(C)** Genomic locations and nucleotide identities of clade-defining SNPs and **(D)** their frequency among SARS-CoV-2 genomes in GISAID.

Identification of recombinant genomes is difficult because the clade structure of SARS-CoV-2 is driven by such a limited number of SNPs. The limited number of SNPs that distinguish certain clades are also often clustered in short regions of the genome, further restricting our ability to reliably identify recombinant genomes. For instance, clades 20B-2 and 20A-2 are primarily distinguishable based on four cdSNPs (25563 and 28881-3), but those four positions span only 3.3 kb of the viral genome. As a result, recombination between strains from clades 20B-2 and 20A-2 throughout the first 80% of the genome would not be detectable. Further, a recombination event that unlinks the nucleotides at positions 25563 and 28881-3 could be parsimoniously explained as a *de novo* T to G mutation at position 25563 in a clade 20A-2 genome or a *de novo* G to T mutation at this position in a clade 20B-2 genome.

Nevertheless, there are many circumstances where detection of recombination should be feasible and where recombination could explain cdSNP patterns more parsimoniously than *de novo* mutation. In particular, all major clades are most strongly differentiated from each other based on 11 SNPs that are distributed throughout the genome (sites 241, 3037, 8782, 11083, 14408, 23403, 25563, 28144, 28881-3). Rearrangement of multiple of these clade-defining markers would be among the strongest indication of recombination between SARS-CoV-2 strains. For instance, the triple mutation GGG to AAC at positions 28881-3 is uniquely found in clade 20B, and would be a strong marker of recombination between clade 20B and clades 19A and 19B when combined with positions 241, 3037, 14408, and 23403.

### Methods that rely on phylogenetically uninformative SNPs are prone to error in detecting recombination

The first report of recombination among SARS-CoV-2 genomes was a correspondence article^6^, prepared at a time when there were 85 SARS-CoV-2 genomes in GISAID. This article argued that the distribution of 4 SNPs in those early genomes could be explained by multiple recombination events. With such little phylogenetic information, it is difficult to evaluate the strength of these claims. However, three of the four sites on which they base their argument are, by our analysis, associated with discrete clades and their distribution is readily explained via a clonal pattern of descent. The remaining site, C29095T is not one of the cdSNPs we identified, but is a low frequency allele that is found in multiple clades^19^and is thus very likely a homoplasy. Consequently, inference of recombination in this study may have been biased by a low sample number and the use of phylogenetically uninformative SNPs.

Similarly, an article currently available on the *bioRxiv* preprint server used a three-way sequence comparison tool, Recombination Analysis Program (RAPR)^20^, to detect recombination between genomes isolated from the same geographic location^7^. Since RAPR incorporates all SNPs equally, regardless of how phylogenetically informative they are, RAPR has the potential to falsely identify putative recombinants and their predicted parent sequences. For example, RAPR identified the genome EPI_ISL_418869 as a recombination product of parent sequences EPI_ISL_422974 and EPI_ISL_422983, with an estimated significance of p=0.0002 (Fig. 2A). However, there are 7 cdSNPs present in the putative recombinant that correspond to neither of the two parent sequences’ cdSNPs (Fig. 2B), demonstrating that recombination between the two parent sequences is exceedingly unlikely to have generated sequence EPI_ISL_418869. Further, the cdSNP profile of EPI_ISL_418869 is a perfect match to clade 19B-1, indicating that it is a typical sequence of that clade (Fig. 2B).

**Figure 2.**
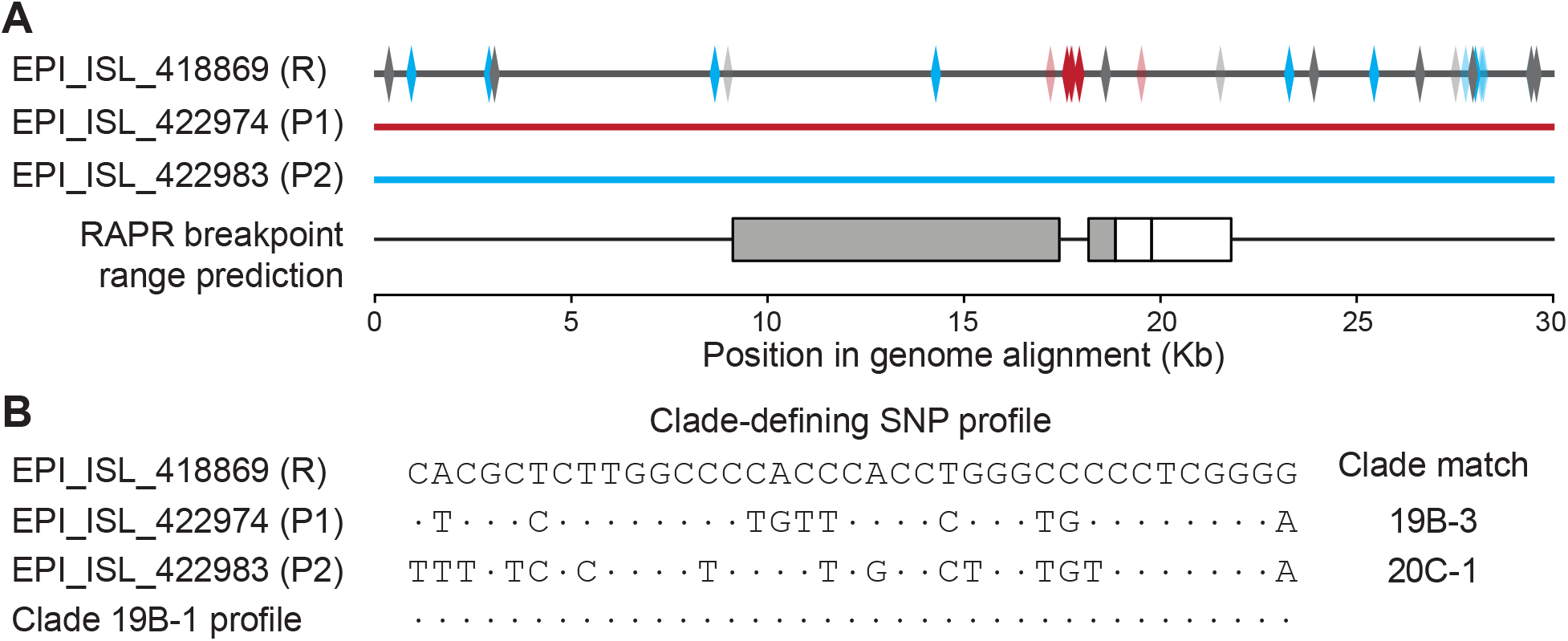
Recombination Analysis Program (RAPR) can falsely identify SARS-CoV-2 sequences as recombinants. **(A)** Three genomes from Washington State, USA, that RAPR identifies as potentially parent and recombinant sequences (p=0.0002). Boxes in the RAPR breakpoint range prediction indicate the ranges where the breakpoint intervals most likely fall; grey boxes indicate the statistically significant range. SNPs in the predicted recombinant sequence (R) that match each parent sequence are shown in blue (P1) and red (P2), while SNPs that do not match either parent sequence are shown in grey. SNPs from the 37 clade-defining SNPs are solid colors (blue, red, and grey), and low-frequency SNPs are partially translucent. **(B)** The cdSNP profile of each sequence and the clade which these profiles match perfectly to are shown. Nucleotides that match the putative recombinant are denoted with a dot. Clade matches have no differences across any of the 37 cdSNPs.

### Clade-defining SNPs support recombination in 1175 genomes

Having identified the SNPs that shape the clade structure of SARS-CoV-2, we next aimed to determine if any of the genomes available on GISAID have combinations of cdSNPs that can be most parsimoniously explained by recombination. In total, we screened 300592 unique genomes and identified 1175 putative recombinant genotypes that have at least 2 cdSNPs supporting recombination (Fig. 3). These putative recombinants were detected between one and 250 times in GISAID and isolated from 57 different countries. These putative recombinants include 847 genomes that harbor the D614G substitution, 8 with N501Y, and 5 with E484K. Notably, two N501Y recombinant genotypes appear to be spreading, having been sampled multiple times in South Africa and in multiple provinces in Belgium. A further 101 genotypes were also detected more than once, but only 14 were detected more than 10 times. Thus, instances of onward transmission generally appear to be limited in their extent.

**Figure 3.**
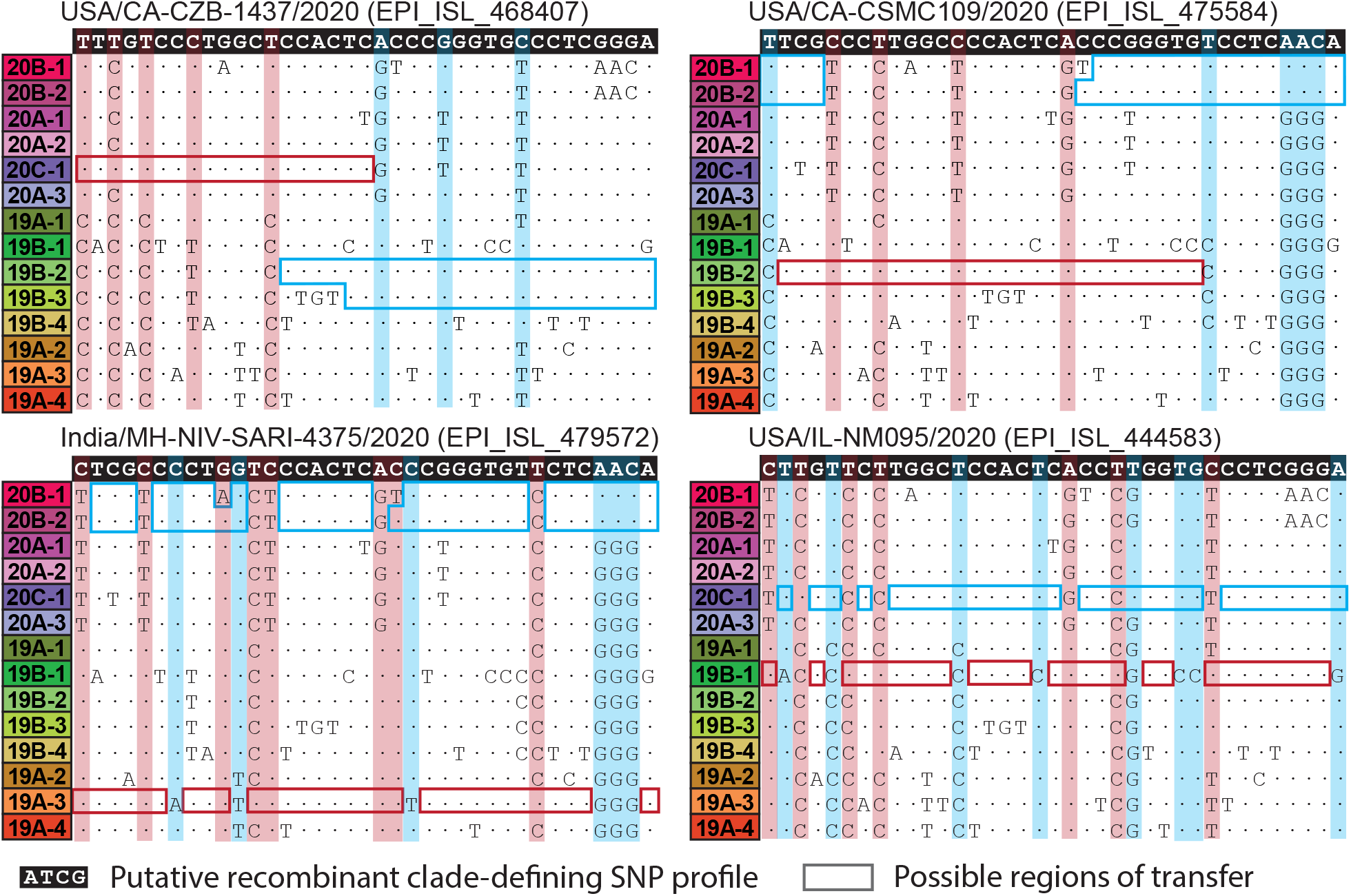
Examples of putative recombinant genomes. Possible parent clades of putative recombinants are identified by screening pairwise combinations of clades for cdSNPs that support recombination without any conflicting nucleotides. The clade-defining SNP profile of each putative recombinant sequence (white text with black background) is compared with the profiles of all 14 clades. Nucleotides that match the putative recombinant are denoted with a dot. Regions boxed in blue and red show potential parental clades, with left and right boundaries indicating potential transfer regions. For some sequences, multiple pairs of parental clades are possible. The minimum number of recombination breakpoints required to explain each genotype ranges from 1 to 10. Clade-defining SNPs that support recombination are highlighted with vertical blue and red windows. Genomes with ambiguous nucleotides at cdSNP positions and those where no possible pair of parent clades can explain the genotype through recombination were excluded.

However, two recombinants appear to be more widely circulating. The most prevalent recombinant was first detected in Kentucky, USA in early August 2020 (EPI_ISL_845666) and its lineage currently comprises 250 GISAID samples spread across 33 additional states and four countries (England, Singapore, Japan, and Canada). The second most abundant recombinant was first detected in England in late September (EPI_ISL_577075) and its lineage currently comprises 100 GISAID samples, predominantly from the US. It is now detectible in 19 different states and three countries. Both recombinant genotypes have D614G substitutions, but no other substitutions in the spike protein or in other proteins that are currently suspected of altering transmission characteristics. As such, we have no reason to believe that these recombinants have altered transmissibility or virulence. Nevertheless, these genotypes mark the first instances of widespread transmission of SARS-CoV-2 recombinants.

Our baseline analysis identifies genomes with at least 2 cdSNPs supporting recombination and any number of recombination breakpoints. We chose not to consider sequences with only one atypical cdSNP, since this may be more parsimoniously explained by *de novo* mutation. The minimum number of breakpoints necessary to generate the putative recombinant genotypes we detected ranges between 1 and 10, but 88% of the genotypes could be explained by between 1 and 4 breakpoints (Fig. 4A). This range in the number of breakpoints is consistent with breakpoint patterns observed in experimental coronavirus co-infections^21^. Increasing stringency to ≥3 cdNPs supporting recombination or limiting the number of allowable breakpoints necessary to explain a genotype via recombination lowers the number of genomes identified (Fig. 3B), but does not impact our conclusion that recombinant genomes are present but currently rare overall.

**Figure 4.**
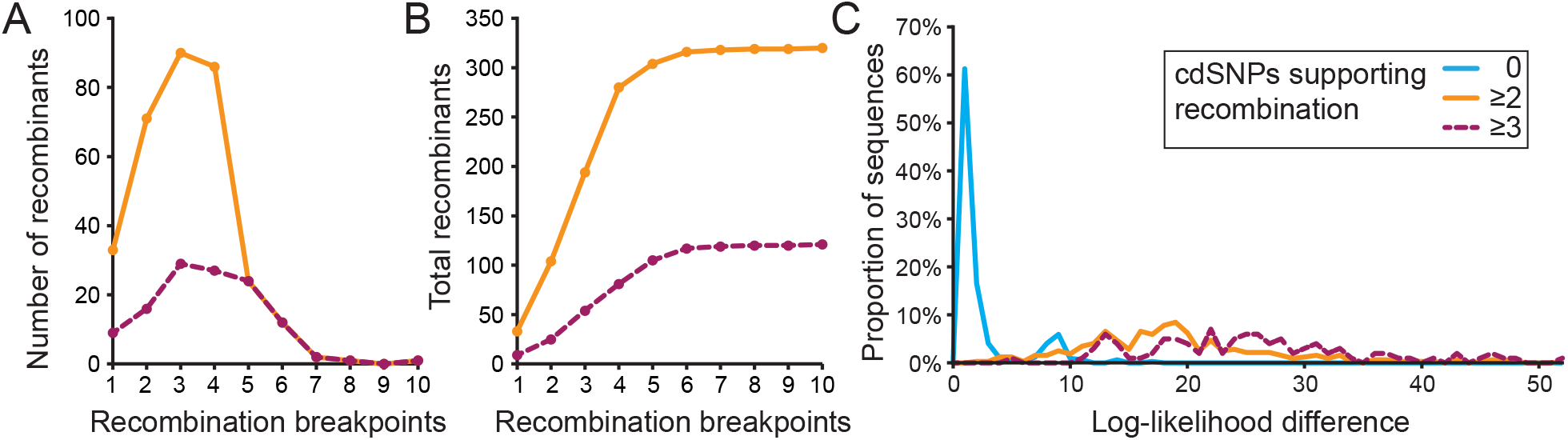
Recombination breakpoint distribution among putative recombinants. **(A)** Histogram showing the distribution for the number of recombination breakpoints needed to explain the cdSNP profile of a recombinant, for the 320 unique recombinant genotypes identified. Yellow line shows the distribution for the subset of recombinants that are supported by 2 or more cdSNPs. Red line shows this distribution for recombinants that are supported by 3 or more cdSNPs. **(B)** Total number of recombinant genomes detected at each breakpoint cutoff by the number with ≥2 (yellow) or ≥3 (red) cdSNPs supporting recombination. **(C)** Log-likelihood difference of mapping the two genome subsets of an identified recombinant and its full genome. Log-likelihood differences are shown separately for those recombinants supported by 2 cdSNPs and those recombinants supported by 3+ cdSNPs. The null expectation, generated from a set of non-recombinant genomes, is also shown.

### Phylogenetic placement analysis provides statistical support for the cdSNP screening approach

Instances of recombination are most strongly supported by significant phylogenetic incongruence, with genome subsets derived from one parent falling phylogenetically in a different part of the phylogeny than genome subsets derived from the other parent. To assess whether our lightweight screening approach delivers putative recombinants that show significant phylogenetic incongruence between their parentally-derived genome subsets, we conducted a phylogenetic placement analysis. This analysis determined whether the likelihood associated with the phylogenetic placement of the two genome subsets significantly exceeded the likelihood associated with the phylogenetic placement of the full genome. To perform this statistical assessment, we first divided each recombinant genome into genome subsets that correspond to the predicted parent clades. To do this, we identified the approximate locations of recombination breakpoints as the midpoint between cdSNPs that support recombination, and then used these locations to subdivide the alignment to create two complementary genome subset sequences. These genome subset sequences each contained all genomic regions derived from one but not the other of the two predicted parent clades. We then mapped the full length and two genome subset sequences to a maximum likelihood reference tree using pplacer^22^ and measured the extent to which subdividing the genome increases the overall likelihood of mapping (Fig. 4C). In the case of recombinants, the overall likelihood associated with mapping of the genome subset sequences will precipitously exceed the likelihood associated with mapping of the full-length sequence. Importantly, these alignments include all polymorphic sites, not just the 37 cdSNPs. For the recombinants supported by ≥2 cdSNPs, mapping 315 out of the 320 unique identified genomes as the two genome subsets resulted in significantly higher log-likelihoods than mapping the full genome (p>0.05). Similarly, mapping all 100 of the identified genomes supported by ≥3 cdSNPs resulted in significantly higher log-likelihoods than mapping the full genome.

To confirm that these log-likelihood differences statistically support the identification of a recombinant, we further generated a log-likelihood difference null expectation from non-recombinant genomes. To generate this null expectation, we sampled 320 non-recombinant genomes and cut them according to the same pattern of breakpoints as the 320 unique putative recombinant genomes. Using pplacer^22^, we then calculated the log-likelihoods of mapping the two genome subsets versus the full genome and plotted their difference as the null expectation (Fig 4C). As expected, the log-likelihood differences from the non-recombinants were considerably smaller than those of the recombinants, indicating that subdividing genomes according to recombinant breakpoints substantially improves mapping among recombinant genomes with either ≥2 or ≥3 cdSNPs supporting recombination, but not among non-recombinant genomes (Fig. 4C). Interestingly, we find weak bimodality in the null expectation, which could reflect low levels of within-clade recombination not detectible by our cdSNP based method. Accordingly, for all remaining analyses, unless otherwise stated, we included all putative recombinants with ≥2 cdSNPs supporting recombination regardless of the number of breakpoints detected.

Visualizing how genome subsets cluster with existing clades on the reference tree following phylogenetic placement further supports recombination. In all detected cases of recombination, the subdivided recombinant genomes clustered closely with genomes from the predicted parent clades (Fig. 5). In cases where multiple predicted parent clades were possible based solely on the 37 cdSNPs (Fig. 5 circle, triangle, and square), phylogenetic placement analysis showed that the recombinant sequence clustered preferentially at the root of the possible parent clades, highlighting that in certain instances there is insufficient information to determine the precise parent clade (e.g. 20A-1 vs. 20A-2 for triangle and square blue regions).

**Figure 5.**
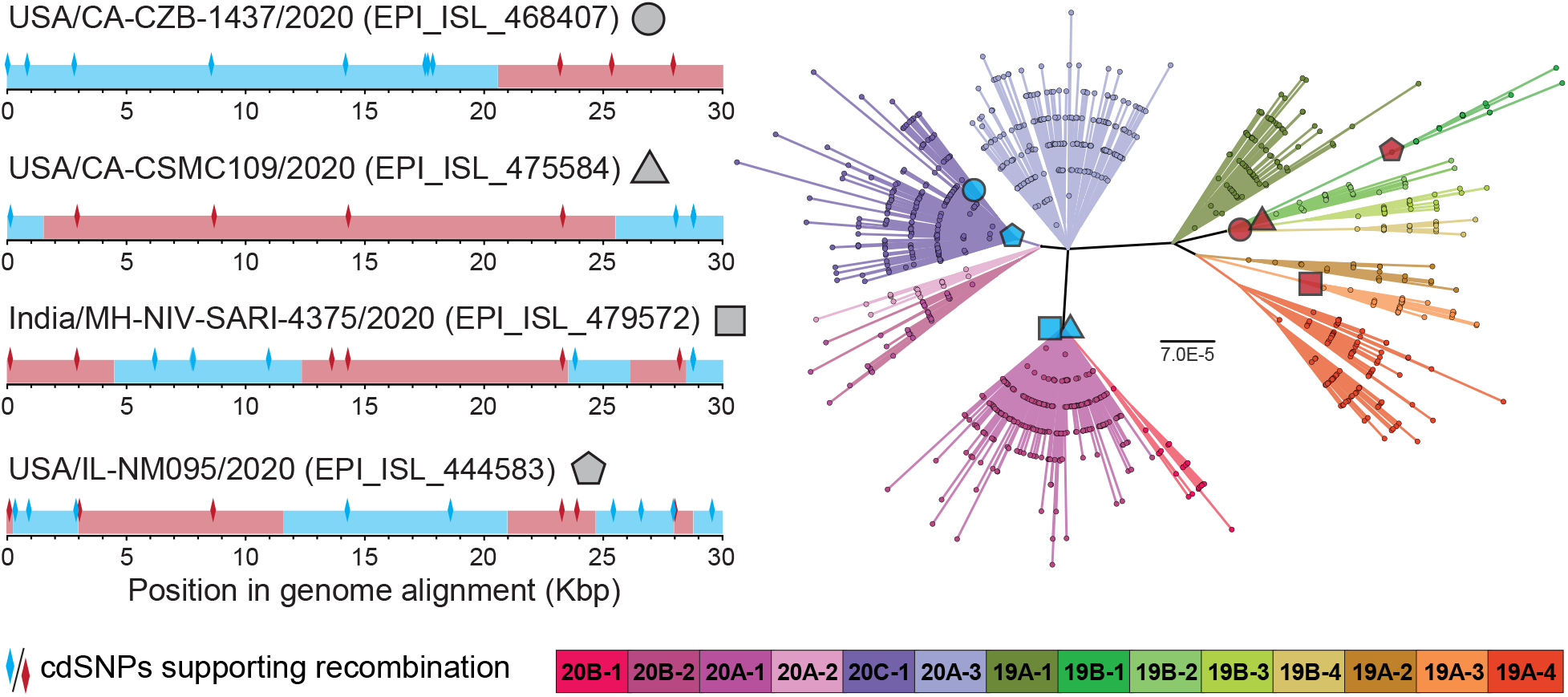
Phylogenetic placement analysis of recombinant genome regions supports parental clade prediction. Genomes were mapped to a maximum likelihood reference tree (inferred under the General Time Reversible model with invariant sites) using pplacer^22^. All four putative recombinant genomes shown here have at least 2 cdSNPs supporting recombination. The putative recombinant genomes differ in the number of recombination breakpoints necessary to explain their cdSNP profiles. The locations of cdSNPs mapping to each parent clade are shown with blue or red diamonds. cdSNPs that are shared by both predicted parent clades are not shown.

### Plausibility of recombinants based on geographic considerations

We next sought to assess, based on geographic considerations, the plausibility of transfer between the predicted parental clades to generate the observed recombinants. We counted all non-recombinant genomes that map to the 14 clades that were sequenced in the location of exposure in the two weeks prior to the collection date of the first instance of each recombinant (Fig. 6). Of the 314 unique putative recombinant genotypes with complete isolation metadata, 181 (58%) were first isolated from individuals who were exposed to SARS-CoV-2 in a country where at least 100 genomes had been sequenced in the two weeks prior to isolation. Of these 181 putative recombinants, 80% had both of the predicted parent clades isolated in the two weeks prior to isolation, demonstrating that the feasibility of co-infection was high.

**Figure 6.**
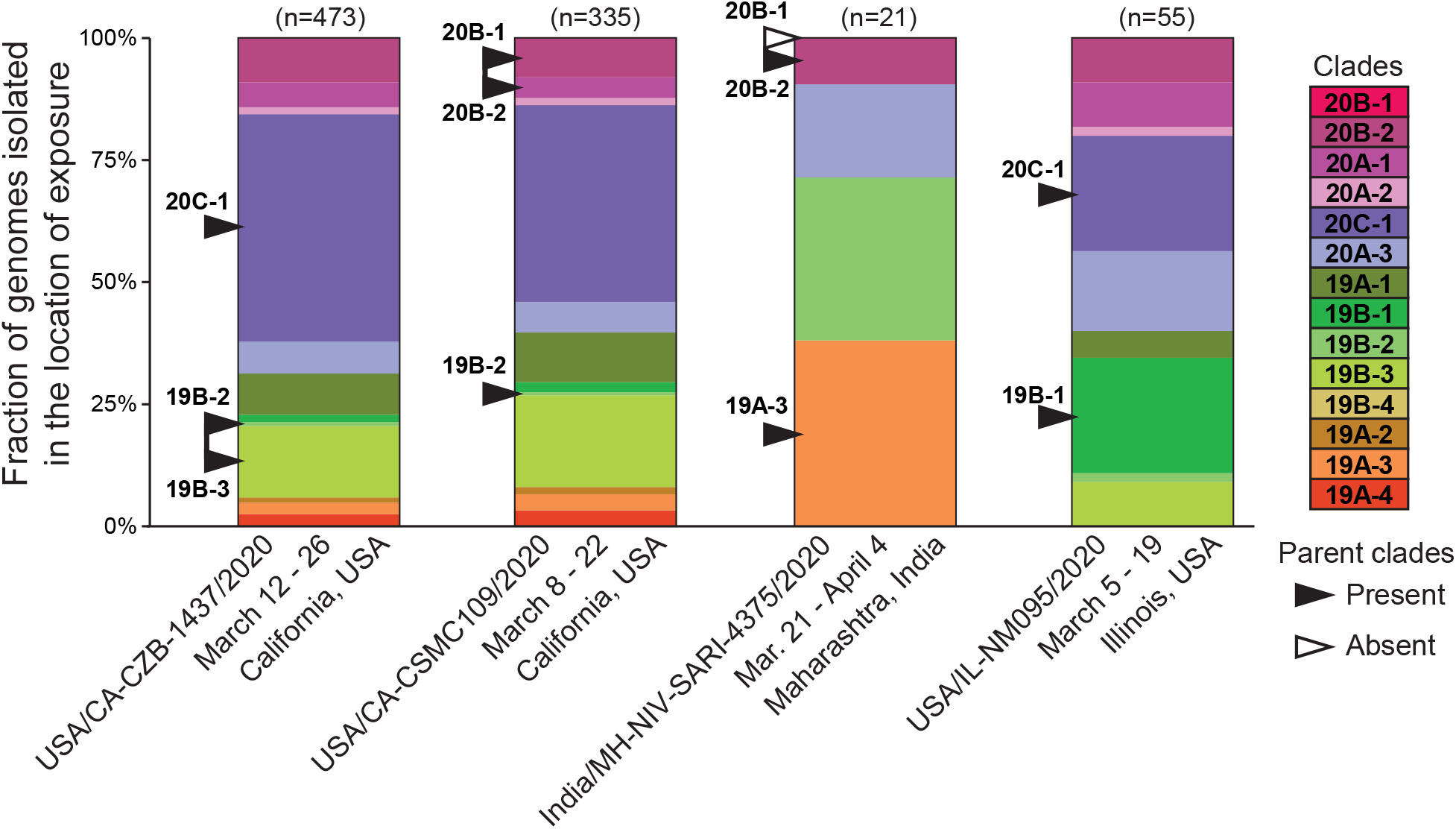
Predicted parental clades of recombinant viruses are frequently detected circulating in geographic locations of exposure prior to the collection dates of recombinants. Predicted parent clades for each listed recombinant genome are shown with arrows. Where multiple parent clades are predicted, both are shown and connected.

It is possible that the chimeric sequences we identified are artefacts of cross-sample contamination, co-infection, or issues related to sample processing and library preparation. Ideally, we would examine the raw reads of each recombinant to determine the likelihood of these alternatives to true recombination. Unfortunately, the raw sequencing reads are not available for the vast majority of genomes deposited on GISAID. Of the recombinants we identified, only 85 could be linked to accession numbers on the NCBI Sequence Read Archive. Sixty-six of these genomes (78%) have high quality reads with no minor alleles detectable at any of the cdSNP sites, supporting that these genomes are recombinant. Only nine genomes with high coverage (11%) are polymorphic at ≥2 cdSNP sites. These genomes are consistent with either co-infection or contamination. An additional 18 genomes (21%) have low coverage at one or more cdSNP sites that are polymorphic, indicating low sequencing coverage could have impacted the quality of the final assemblies. However, all but five of these low-coverage genomes have the same cdSNP profile as high-quality/coverage genomes, suggesting many of these genomes could still be true recombinants. Overall, among those putative recombinant genomes for which raw sequencing data are publicly available, the majority are well-supported and unlikely to be due to co-infection or sequencing artefacts.

### No evidence for hotspots of recombination in the SARS-CoV-2 genome

Since we detected a substantial number of putative recombinants, we next sought to use these sequences to ask whether recombination hotspots in the SARS-CoV-2 genome may be apparent. The limited genome diversity restricts our ability to identify discrete regions where recombination breakpoints occur. Instead, we identified ranges where breakpoints could have occurred based on the location of cdSNPs (Fig 7A). Next, we generated a simulated dataset that was designed to represent the null expectation that recombination breakpoints occur randomly across the genome. Simulated recombinant genomes were generated such that they have the same distribution of breakpoints per genome as the GISAID recombinants, but the locations of those breakpoints in the genome were random. Each simulated genome was generated by taking two random parent clades, picking random locations in the genome to create breakpoints, then assigning the identity of cdSNPs according to the location of those breakpoints and the parent clades used. Importantly, the probability of selecting each parent clade was proportional to its world-wide abundance in GISAID and the resulting simulated genomes were then screened using the same scripts used to screen GISAID sequences to ensure they passed the same criteria. Comparing the observed and simulated recombinant breakpoint ranges did not reveal a substantial enrichment anywhere in the genome (Fig. 7B), regardless of their proximity to genes or transcription-regulatory sequences (Fig. 7C). Thus, our results suggest that breakpoints occur randomly and are not substantially structured by genome composition or biological processes. Alternatively, our currently limited dataset of recombinants may simply not have the power to detect low levels of recombination hotspot activity.

**Figure 7.**
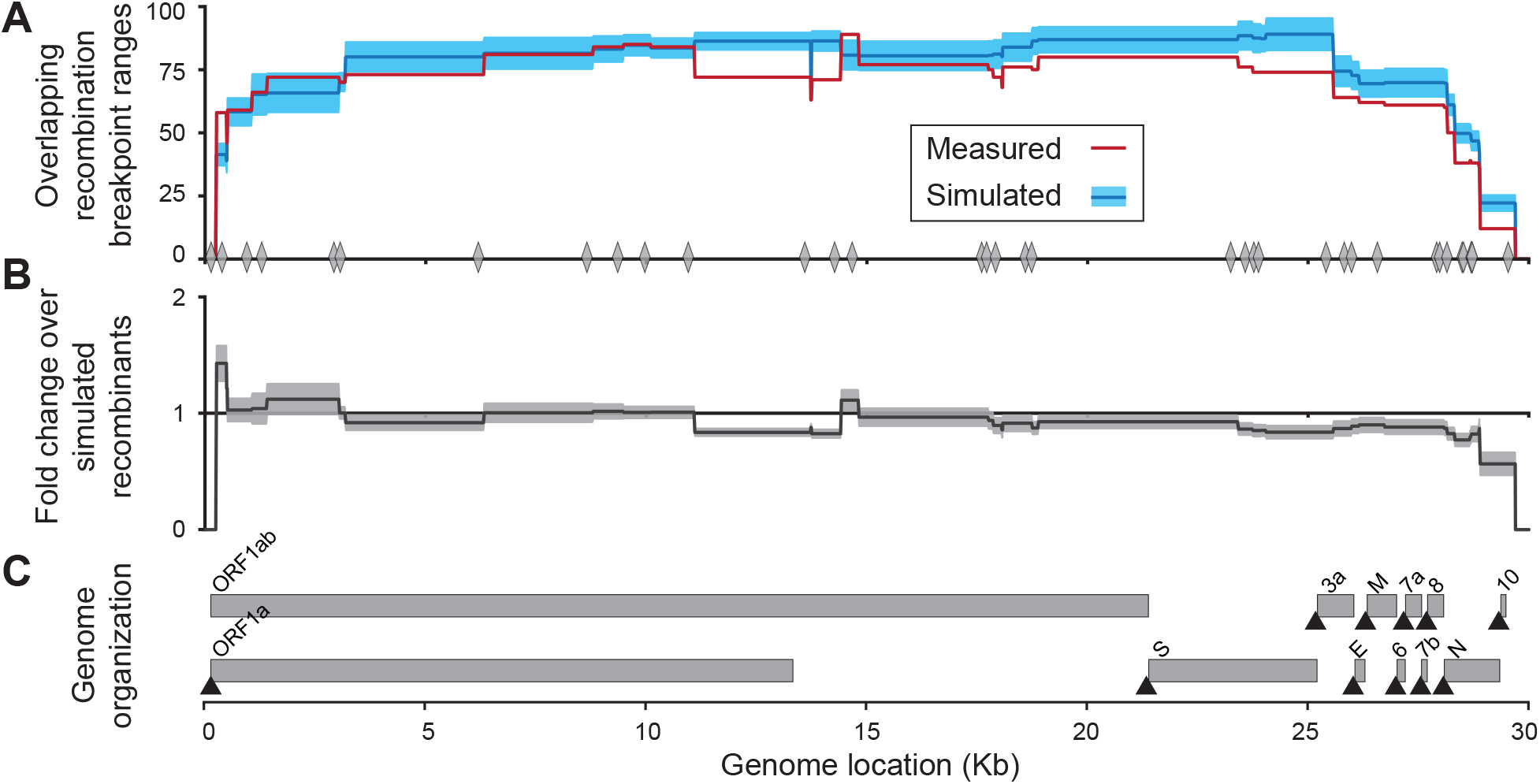
Identification of hotspots for recombination breakpoints. **(A)** The number of overlapping recombination breakpoint ranges at each site in the SARS-CoV-2 genome from the 320 unique recombinant genotypes (red) and an equal number of simulated recombinant genomes (blue). 95% confidence intervals are shown in shaded blue. The location of the 37 clade-defining SNPs are shown along the X-axis with grey diamonds. **(B)** Fold change in the number of overlapping breakpoint ranges of the observed recombinants over simulated recombinants. **(C)** Location of the open reading frames and transcription-regulatory sequences (black triangles) in the SARS-CoV-2 genome.

### At most 5% of circulating viruses are recombinants

Through our analysis we identified 1175 putatively recombinant genomes out of 537360, corresponding to a frequency of recombinants of 0.2%. It is important to note that this detected frequency of recombinants is a lower bound on the frequency of recombinants in circulation. This is because some recombinants will go undetected because they involve two parent clades that have highly similar cdSNP profiles. Indeed, the probability of detecting a recombinant depends on the number of SNPs that differentiate its parent clades (Fig. 8A). As such, the relative abundance of the 14 clades in a given geographic location (along with sampling intensity) determines the chance that a recombinant genome is detected if it is in circulation. By considering these factors impacting recombination detection, we were able to estimate a ceiling on the proportion of circulating viruses that are recombinant. We estimated this ceiling for the USA and for the United Kingdom separately, due to markedly different relative abundances of the 14 clades in these two regions (Fig. 8B). We chose these geographic locations due to their large sequencing efforts relative to other locations. For these two countries, we estimate at most 1-5% of circulating viruses were recombinant (Fig. 8C).

**Figure 8.**
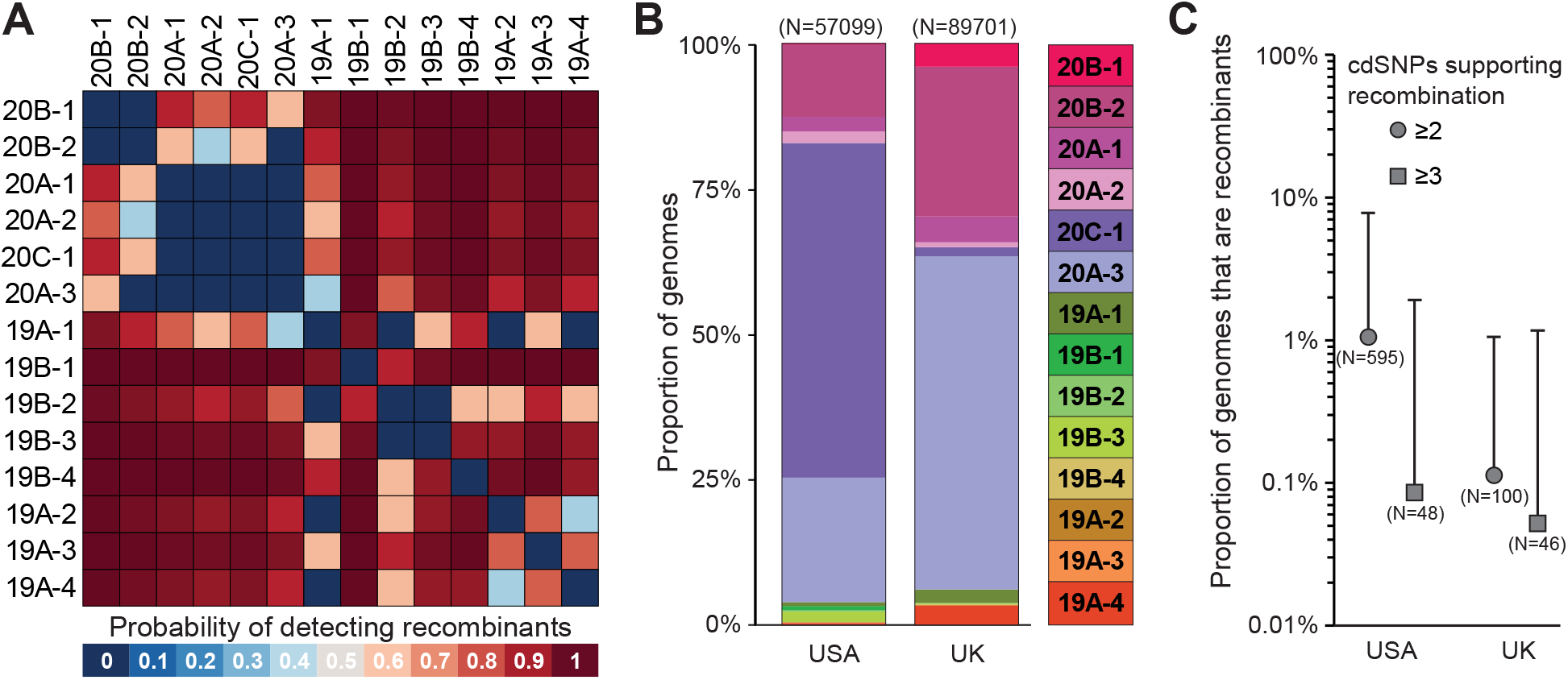
Estimation of the maximum proportion of the circulating virus population that is recombinant. **(A)** The probability of detecting a recombinant from pairwise combinations of parent clades. Here, we required at least 2 cdSNPs to support recombination. **(B)** The relative abundance of each clade in GISAID from the USA and United Kingdom, calculated from SARS-CoV-2 sequences collected prior to February 16, 2021. **(C)** Proportion of genomes that are recombinant in the USA and UK. The top of the error bar shows the maximum estimated proportion of recombinants after accounting for local clade frequencies. Circle and square datapoints show the proportion of genomes that are recombinant with at least 2 and 3 cdSNPs supporting recombination, respectively. The total number of recombinant genotypes used in each analysis are shown in parentheses.

### Comparison with Bolotie, an alternative method for detecting SARS-CoV-2 recombination

A preprint posted on *biorxiv* recently also focuses on the identification of SARS-CoV-2 recombinant genotypes^23^. The method presented, called Bolotie, outperforms other methods, such as the Recombination Analysis Program (RAPR)^20^ used by Korber *et al*.^7,24^ in their preprint on *medrxiv*, in detecting recombinant genotypes for two reasons: it incorporates only those nucleotide variants present in at least 100 genomes and it weighs the influence of each SNP in supporting recombination by the strength of association that the SNP has with each clade. Owing to similarities between this approach and our own, there is substantial overlap in our results. Although the authors of Bolotie do not provide a list of the 225 recombinant genomes they detected from 87695 genomes (0.3%), we ultimately detect a similar proportion of putatively recombinant genomes with our method. The authors explicitly reference eight putative recombinants in their main text, and six overlap with sequences we identified as recombinants (EPI_ISL_489588, EPI_ISL_439137, EPI_ISL_468407, EPI_ISL_417420, EPI_ISL_452334, EPI_ISL_475584). One of the remaining sequences, EPI_ISL_510303, was identified by Bolotie as a recombinant, and would have been flagged as putatively recombinant by our analysis, but was excluded because one of the 37 cdSNPs was identified as an ambiguous nucleotide (site 2919). Ultimately, Bolotie and the method we present here provide complementary evidence of recombination among circulating SARS-CoV-2 strains. Future studies that systematically evaluate the strengths and weaknesses of each method will be valuable to identify the most sensitive and reliable approaches for detecting recombination in SARS-CoV-2.

### Continued surveillance

Since we downloaded the 537360 complete SARS-CoV-2 genomes available on GISAID for the analysis in this paper on February 16, 2021, an additional 160000 genomes have been deposited as of March 8, 2021. As this number continues to grow, continued surveillance will greatly benefit from a fast and lightweight method for screening new genomes for recombinant sequences. While we are able to use previously developed tools like RDP4^25^ to confirm the genomes we detected are recombinants, most tools are not well suited to the combination of large database sizes and low levels of genome-wide diversity, both of which characterize current SARS-CoV-2 sequence datasets. Although our approach was successful in screening over half a million genomes, its efficiency and user accessibility are limited by the large resource demands needed to generate whole genome alignments. With this need in mind, we developed a lightweight version of our pipeline that requires only blastn^26^ and python to generate local alignments to identify cdSNPs. This method, termed cladeSNP-blast, can screen 10K genomes in 15 minutes on a 3.5 GHz single-core CPU with no loss in specificity. The code is available on GitHub (https://github.com/davevanins/Sars-CoV-2_CladeSNP).

## Discussion

The small number of polymorphic sites in the SARS-CoV-2 genome that are phylogenetically informative means detecting recombinant genomes is difficult, and highly dependent on the identity of the parent clades. By identifying the nucleotide changes that underpin the clonal phylogeny of SARS-CoV-2, we established criteria for identifying putative recombinant genomes, and for evaluating their plausibility. We analyzed 537360 SARS-CoV-2 genomes from GISAID and found 1175 viral sequences that contain evidence of recombination. These genomes have rearrangements of multiple cdSNPs that support recombination between clades and are typically isolated in countries where the predicted parent clades are prevalent. Although the number of recombinants depends heavily on the criteria used to distinguish true recombinants from *de novo* mutation or other sources of error, we estimate that the number of circulating recombinant viruses remains low (<5%).

While we were able to identify more than 1000 putatively recombinant SARS-CoV-2 genomes, they represent an extremely small fraction of the genomes available on GISAID (0.2%). This observation supports reports that have found no evidence of widespread recombination among SARS-CoV-2 genomes^9–11^. Indeed, examining the pattern of cdSNPs suggests none of the 14 clades we identified emerged through recombination (Fig. 1). The only site that does not strictly follow the pattern of vertical descent is C14805T, which occurs in both clades 19A-4 and 19B-4. However, none of the seven other cdSNPs that differentiate these clades support recombination, suggesting C14805T is a homoplastic site.

The low frequency of recombinant sequences in GISAID could be due to multiple factors. First, due to the limited genetic diversity of SARS-CoV-2 at this point in time, a large fraction of viral recombinants may not be detectable. This is particularly the case if the parental clades share a large number of cdSNPs (Fig. 8A). However, this cannot be the sole factor, given that our ceiling estimates that take into consideration the inability to detect recombinants rising from pairs of parent clades lie between 1% and 5%. Second, recombinant genomes may be rare because coinfections only rarely occur. Coinfection may be infrequent for SARS-CoV-2 given the acute nature of the infection and that some geographic regions (but not others) have managed to keep the level of virus circulation low. Third, recombinant genomes could be rare because they are transmitted infrequently. For instance, when coinfections do occur, recombinant genomes may evolve late in the infection, resulting in rare onward transmission.

It is likely that a portion of the putative recombinants we identify here are non-recombinant sequences that reflect issues with library preparation, sequence depth limitations, and coinfection or contamination. Our analysis nevertheless suggests the majority of the putative recombinants we identified are based on high-quality sequences with multiple independent lines of evidence supporting the feasibility of recombination. In particular, 105 of the unique putatively recombinant genotypes occurred more than once in GISAID, and the majority of these genotypes were sequenced multiple times by different laboratories in the same country or independently isolated in other countries. These observations suggest that many of the sequences we identified cannot be accounted for by sequencing artefacts. Although the number of recombinant genomes in GISAID is difficult to define precisely, small variations in the total number do not substantially affect our estimates of the maximum proportion of recombinant viruses that are currently circulating.

Since one of the limiting factors in identifying recombinant genotypes is the small number of phylogenetically informative sites, it is tempting to assume that it will become a progressively easier process as mutations continue to accumulate. However, the ability to detect recombinants depends on the identities of the circulating clades and these may change over time, in part due to viral adaptation. For example, the probability of detecting a recombinant is highest between clade 19 and 20 parents, and lowest between clade 20 parents (Fig. 8). Over the last 4 months, clade 19 viruses have become progressively rarer while the D614G harboring clade 20 viruses have disproportionately driven waves of infection around the world. As a result, recombinants have become more difficult to detect than when clade 19 and 20 viruses co-circulated more uniformly. Similarly, recent waves of infection driven by three clade 20 N501Y bearing lineages have further reduced our ability to detect recombinant genomes. Accordingly, we expect that the identification of SARS-CoV-2 recombinant genomes may continue to be difficult as novel adaptive mutations continue to drive new waves of infection.

Ultimately, our results suggest that recombination between SARS-CoV-2 strains is occurring, but these chimeric genotypes remain rare. Although we identify a small number of recombinant genotypes that are actively circulating, we have no reason to expect that these lineages – or any other recombinants identified here – have increased transmissibility or virulence. Yet, as novel mutations that influence transmissibility or threaten to limit the efficacy of vaccines continue to evolve and spread, the potential for recombination to facilitate merging these mutations into a single background will continue to increase. Given our finding that recombination is already occurring in SARS-CoV-2, surveillance efforts and real-time analyses to detect recombinants, such as the one here, should be sustained to monitor the circulation and potential spread of high-fitness recombinant genotypes.

## Materials and Methods

### Genome quality filtering and alignment

Genomes were downloaded from the GISAID genome databases^1^, and filtered to exclude low quality sequences. All genomes were trimmed relative to positions 118 and 29740 in the NCBI reference sequence (accession NC_045512) because these regions are inconsistently assembled between genomes and increase resource demands and uncertainty in following steps. To trim genomes at these locations precisely prior to whole genome alignment, the 118 and 29740 sites were identified using BLASTn^26^. After trimming, genomes with less than 1% Ns and a final length greater than 29,610 bp and less than 29,660 bp were included in further analysis. Genomes were aligned to the NCBI reference sequence genome using MAFFT v7.464^27^, using the option “keeplength” to exclude any insertions not present in the reference sequence. Excluding insertions reduced resource limitations and enables parallelization since all genomes are aligned to the same reference. Only samples collected prior to March 1, 2020 were considered.

### Identifying clade-defining SNPs in SARS-CoV-2 genomes

Clades were identified as monophyletic groups with at least 50% bootstrap support within a maximum likelihood phylogenetic tree built from 6536 unique high quality genome sequences using PhyMLv3.1^28^. Reference genomes were picked by clustering all genomes available on GISAID from February 1 to July 1, 2020, based on their pairwise distances on a neighbor-joining tree constructed from a whole genome alignment. Representative strains were picked to minimize redundancy while maximizing the total sampled diversity. Accession numbers of these reference genomes are available on our GitHub repository (https://github.com/davevanins/Sars-CoV-2_CladeSNP). Clades were named according to the Nextstrain clade to which they belong, with added suffixes of ‘-1’, ‘-2’, etc. to denote clades at finer resolution than those available under the Nextstrain system. Clade-specific SNPs (cdSNPs) were subsequently identified as SNPs that are present in >95% of all members of a clade while >95% of the members in remaining clades had another nucleotide at that position. Recombinant genomes were identified by comparing the cdSNP profiles of each query sequence against the profiles of the 14 clades. Any sequences with at least two cdSNP differences from the nearest clade cdSNP profile were screened to determine that the genotype could be explained by recombination between two parent clades. This minimum distance to the nearest clade cdSNP profile represents the minimum number of cdSNPs supporting recombination. All genomes with two possible parent clades based on cdSNPs that pass all other quality thresholds were considered putatively recombinant. Recombinant genomes with ≥2 and ≥3 cdSNPs supporting recombination are frequently analyzed separately to represent different levels of stringency in identifying putative recombinants. The number of breakpoints in a putative recombinant was estimated as the minimum number of breakpoints required to explain the parental origin of the genome’s cdSNPs. In total, 537360 genomes were screened. The list of their GISAID accession numbers can be found on the GitHub repository for the analyses contained within this manuscript (https://github.com/davevanins/Sars-CoV-2_CladeSNP).

### Phylogenetic placement analysis

Phylogenetic placement was performed to provide statistical support for recombination. After predicting the parent clades that minimize the number of cdSNP differences and recombination breakpoints, putative recombinant genomes were subdivided according to the midpoints of the recombination breakpoint ranges. These genome subsets were then mapped onto the maximum likelihood reference phylogenetic tree using pplacer^22^, which provides their log-likelihoods of placement along particular branches. Placement on the reference tree was visualized using Interactive Tree of Life (iTOL)^29^. Significance of mapping was determined by the log-likelihood difference between the combined log-likelihood of mapping both genome subsets and the log-likelihood of placement for the full-length genome. Recombinant genomes with ≥2 and ≥3 cdSNPs supporting recombination were compared to a null distribution from non-recombinant sequences. The null distribution was generated by sub-dividing the genomes of 1175 random non-recombinant sequences according to the breakpoints inferred for the observed recombinants, sampled without replacement.

### Quantifying recombination breakpoint frequency

Recombination breakpoint frequency enrichment was quantified by comparing the number of overlapping breakpoint ranges in the 320 unique recombinant genomes with ranges derived from simulated recombinant genomes with random breakpoint locations. Breakpoint ranges were defined as the full region between any two cdSNPs that were predicted to come from different predicted parent clades. To simulate recombinant genomes, we picked two random parent clades and picked random locations throughout the genome to place breakpoints. Based on the locations of the random breakpoints and the 37 clade-defining SNPs, we created cdSNP profiles for each simulated genome by first randomly picking which parent clade the very first cdSNP originated from, then assigning the remaining cdSNPs of the recombinant according to the locations of the breakpoints. The probability of picking any one clade was proportional to its abundance in GISAID. Simulated recombinants were generated such that they had the same distribution of breakpoints per genome as the observed recombinants. To do this, recombinants were generated iteratively by creating genomes with one random breakpoint until enough simulated genomes with ≥2 cdSNPs supporting recombination were identified using the same code used to screen GISAID genomes. This process was then repeated for two through ten random breakpoints, where the number of detected breakpoints was required to match the number inserted. Simulated recombinants with redundant cdSNP profiles were discarded. Fold change in the number of overlapping breakpoint ranges in the GISAID recombinants relative to the simulated recombinants was calculated in 10 nucleotide long bins across the genome. Ten separate simulations were performed to calculate 95% confidence intervals.

### Calculating the ceiling on the proportion of circulating virus that is recombinant

To statistically estimate the maximum proportion of recombinant viruses circulating in a population, we first calculated the probability of detecting a recombinant arising from parent clades *i* and *j*, where *i* and *j* take on values between 1 and 14, the number of parent clades identified using a 50% bootstrap support cutoff value. We define *m* as the number of cdSNP sites at which parent clades *i* and *j* differ from one another. We define *T* as the threshold number of cdSNP differences required to identify a recombinant. (In our analyses, we choose either *T* = 2 or *T* = 3.) If *m* < *T*, then the probability of observing a recombinant between these parent clades was set to zero. If *m*>= *T*, the probability of observing a recombinant between two parent clades *i* and *j* was calculated as follows:

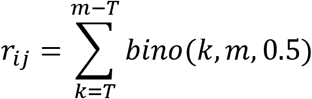

where *bino*(*k*, *m*, 0.5) yields the probability that a recombinant between parent clades *i* and *j* has exactly *k* of the *m* distinguishing SNP sites derived from one parent and the remainder derived from the other parent. This calculation assumes an infinite number of recombinant breakpoints, such that cdSNPs along the genome each have equal probability to belong to parent clade *i* as parent clade *j*. This is particularly useful for estimating the ceiling because it is the least conservative assumption. Fig 8A shows these probabilities for a *T* = 2 cdSNP threshold.

For a given geographic region, we then calculated the frequency of each of the 14 clades from sequences deposited in GISAID prior to February 16, 2021. We denote the frequency of clade *i* as *p*_*i*_. These frequencies are shown in Fig 8B for the USA and UK.

To estimate the ceiling on the proportion of recombinant genomes in circulation, we first calculated the overall probability *D* of a sampled virus being detected as a recombinant under the assumption that a proportion *P_r_* of circulating viruses are recombinant. This overall probability is given by:

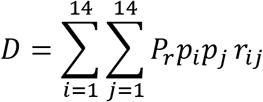

Given *N* sampled genomes from a region, we then calculated the 95% confidence interval for the number of recombinant genomes that would be detected among the number of sampled genomes. This is given by the binomial inverse cumulative distribution function, evaluated at 0.025 and 0.975, with the total number of trials being given by *N* and the probability of success being given by *D*. We determine the ceiling as the value of *P_r_* that yields a lower bound on the 95% confidence interval that exceeds the number of observed recombinants in the data, *n*.

### Assessing assembly quality of recombinant genomes

Raw sequencing reads were downloaded from the NCBI SRA and processed using the BBTools feature “bbduk” to trim contaminating adapter and PhiX sequences and remove reads shorter than 25 nucleotides long, and any reads with poly-a sequences. Reads were then trimmed using trim galore with default settings. The remaining reads were mapped to the SARS-CoV-2 reference genome (accession NC_045512) with “bbmap” with a minimum average quality of 25. Samtools was used to generate a pileup formatted alignment, and assembly quality at the 37 cdSNP locations was assessed by parsing that file using custom python scripts. Assemblies were assessed for the quality and depth of coverage for reads mapped to the 37 cdSNP sites. To determine if co-infection or contamination could have influenced the consensus assembly, indicated by having low-frequency alleles (>10%) at any of the 37 cdSNP positions.

## Code availability

All custom computer code necessary to reproduce our results are available on GitHub (https://github.com/davevanins/Sars-CoV-2_CladeSNP).

## Acknowledgments

This study was supported by NIAID Centers of Excellence for Influenza Research and Surveillance (CEIRS) grant HHSN272201400004C and an Emory University MP3 seed grant. Further support for this study was provided by the US National Institutes of Health National Institute of General Medical Sciences grant 1R01 GM124280-03S1 (supplement). We gratefully acknowledge all of the authors from the originating laboratories responsible for obtaining the specimens and the submitting laboratories where genetic sequence data were generated and shared via the GISAID Initiative, on which this research is based.

## Author Contributions

D.V., K.K., and A.C.L. conceived and designed this study. D.V. performed analyses and wrote the code generated for this study. D.V., K.K., A.C.L., and A.S.N. prepared the manuscript. K.K., and A.C.L. supervised the project and secured funding.

## Competing Interests statement

Authors declare no competing interests.

## Notes

https://www.gisaid.org/

https://github.com/davevanins/Sars-CoV-2_CladeSNP

### Competing Interest Statement

The authors have declared no competing interest.

### Summary of Updates

This revised manuscript contains analysis of >500,000 SARS-CoV-2 genomes from GISAID.org, identifying over 1100 additional recombinant genomes and including new analyses providing statistical support for recombination, measurement of recombination breakpoint frequency enrichment, an estimation for the maximum proportion of circulating viruses that are recombinant, and includes a new lightweight tool for rapidly screening large genome databases for recombinant genomes.

